# Distinct electrophysiological signatures reveal neuronal heterogeneity in the mouse fasciola cinereum

**DOI:** 10.1101/2025.08.13.670170

**Authors:** Salvatore Incontro, Lijun Guo, Miles Dryden, Alicia Garcia-Rivas, Colin Clark, Simra Kazimuddin, Quynh-Anh Nguyen

**Author notes:** Corresponding author: Quynh-Anh Nguyen. These authors contributed equally (Salvatore Incontro, Lijun Guo). **Author Contributions:** S.I., L.G. and Q.A.N. designed research; S.I., L.G., A.G.R, M.D., C.C. and S.K. performed research; S.I., L.G., and M.D. analyzed data; S.I. and Q.A.N. wrote the paper with input from all authors. **Declaration of Interests:** The authors declare no competing interests.

## Abstract

The fasciola cinereum (FC) is a small, conserved hippocampal subregion whose cellular properties remain poorly understood. Anatomically positioned between dorsal CA1 and the third ventricle, the FC receives diverse cortical and subcortical inputs but is often excluded from hippocampal circuit models. Here, we performed *ex vivo* whole-cell patch-clamp recordings in mouse hippocampal slices to define the intrinsic and synaptic properties of FC neurons. We find that FC neurons are electrophysiologically and morphologically distinct from neighboring CA1 and CA2 pyramidal cells, exhibiting reduced intrinsic excitability, delayed spike initiation, and enhanced afterhyperpolarizations. These properties are strongly shaped by potassium conductances, with Kv2 (KCNB) and Kv7 (KCNQ) channels contributing to the suppression of firing. We find that FC neurons span a continuum of intrinsic excitability which is structured by variability in potassium channel–mediated conductances and is inversely coordinated with excitatory synaptic input, such that less excitable neurons receive stronger synaptic drive. Together, these findings establish a mechanistic framework in which intrinsic membrane properties and synaptic input are co-organized across the FC population, providing new insight into how this understudied hippocampal region may contribute to the regulation of neural excitability.

**Key Points:**

- The fasciola cinereum (FC) is a small and poorly understood hippocampal subregion that has recently been implicated in hippocampal related function and seizure activity.
- Here we show that neurons in the FC are morphologically and functionally distinct from cells in neighboring CA1 and CA2 hippocampal subregions, and find substantial physiological diversity of FC neurons, including differences in firing behavior, membrane properties, and intrinsic excitability.
- Pharmacological analyses show that Kv2 and Kv7 potassium channels play a key role in shaping the intrinsic properties of FC neurons.
- Less excitable FC neurons receive stronger excitatory synaptic input, revealing an inverse relationship between intrinsic membrane excitability and synaptic drive.
- These findings identify the FC as a physiologically specialized hippocampal subregion and provide a foundation for understanding its role in hippocampal function and neurological disease.

## Introduction

The hippocampus is classically organized into distinct subregions, including the dentate gyrus (DG), CA3, CA2, and CA1. At its dorsomedial boundary lies the fasciola cinereum (FC), a small and evolutionarily conserved subregion that has received comparatively little attention. The hippocampal formation is classically organized along a well-defined tri-synaptic pathway, with information flowing from the entorhinal cortex to the DG, then to CA3, and subsequently to CA1, before being relayed back to the entorhinal cortex or projected to other cortical and subcortical regions (1). This unidirectional circuit underlies key hippocampal functions such as spatial navigation (2, 3), learning (4, 5), and memory consolidation (6–9). Anatomically, FC is situated at the interface between the third ventricle and dorsal CA1 in rodents. It receives convergent inputs from diverse cortical and subcortical regions, including the entorhinal and perirhinal cortex, and has been found to project primarily to the DG, with additional projections to indusium griseum, retrosplenial cortex, intermediate lateral septum, distal CA1 and septal CA2 (10–16). However, the precise connectivity of the FC remains incompletely resolved, as recent work did not find clear evidence for a prominent projection to the dentate gyrus (11). These distinct and still evolving connectivity patterns point to a unique role of the FC in shaping network interactions within and beyond the hippocampus. Nevertheless, it has long been viewed as a vestigial or transitional zone and excluded from prevailing models of hippocampal circuitry.

Recent work challenges this view, revealing that ablation of FC in rats impaired the acquisition of visual contextual memory (11, 12, 17). In addition, aberrant FC activity can play a causal role in seizure initiation and propagation, as supported by intracranial recordings in human patients showing seizure onset within the FC and optogenetic inhibition studies of FC in a mouse model of temporal lobe epilepsy (TLE) (18). Furthermore, surgical ablation of the FC in a drug-resistant temporal lobe epilepsy patient was associated with seizure relief, suggesting a conserved pathophysiological role across species (18–21). The FC shares several molecular markers with the CA2 hippocampal subregion, which has also been shown to regulate seizures in mouse models of TLE (22–26), suggesting a potential developmental and functional continuum between these regions. Although a recent transcriptomic study highlighted molecular heterogeneity between the FC and CA2 (27), to date there has been no systematic comparison of the electrophysiological and morphological properties between FC and CA2. Understanding how FC neurons differ from their CA2 counterparts is thus essential to elucidate how these hippocampal structures contribute to network modulation and pathological hyperexcitability.

Recent *in vivo* studies in rodent models have reported conflicting results regarding the electrophysiological properties of FC neurons (11, 12, 17). Juxtacellular recordings suggest that FC neurons exhibit firing properties comparable to dorsal CA1 neurons (11), whereas tetrode recordings indicate sparse and delayed firing, consistent with reduced excitability (12). These discrepancies leave unresolved fundamental questions regarding the intrinsic properties of FC neurons and the extent of their heterogeneity. In addition to intrinsic membrane properties, differences in synaptic input may also contribute to shaping neuronal activity across hippocampal subregions. However, whether FC neurons differ from neighboring regions in their excitatory synaptic drive remains unclear. Characterizing both intrinsic excitability and synaptic input is essential to define how FC neurons are positioned functionally within hippocampal circuits.

Here, we performed whole-cell patch-clamp recordings and post hoc morphological analysis in acute mouse hippocampal slices to systematically characterize FC neurons and compare them with CA1 and CA2 pyramidal cells. We find that FC neurons exhibit reduced intrinsic excitability and distinct dendritic architecture relative to neighboring regions. Importantly, we found variation in FC intrinsic excitability which is strongly influenced by potassium channel–mediated conductances. In parallel, FC neurons receive overall enhanced excitatory synaptic input compared to neighboring CA1 pyramidal cells, which varies across the population and is inversely related to intrinsic excitability. Together, these findings identify distinct properties unique to the FC among hippocampal subregions and provide a cellular framework for understanding its potential contribution to hippocampal network dynamics.

## Results

### FC neurons exhibit reduced intrinsic excitability and distinct dendritic architecture

Whole-cell current-clamp recordings were performed in acute hippocampal slices to compare the intrinsic properties of neurons in CA1, CA2, and FC (Fig. 1 A). Representative firing patterns revealed markedly lower excitability in FC neurons compared with CA1 and CA2 pyramidal neurons (Fig. 1 B). Consistent with this observation, the input–output relationship showed a shallower increase in action potential (AP) frequency in FC neurons (Fig. 1 C). FC neurons required larger current injections to reach threshold (rheobase; Fig. 1 D) and exhibited longer interspike intervals (Fig. 1 E), indicating reduced firing capability. These differences were accompanied by distinct intrinsic membrane properties. FC neurons displayed more hyperpolarized resting membrane potentials (Fig. 1 G), higher AP thresholds (Fig. 1 H), and smaller AP amplitudes (Fig. 1 I). Subthreshold and passive properties also differed across regions: FC neurons exhibited smaller hyperpolarization-activated sag responses (Fig. A1 B), larger medium afterhyperpolarization (mAHP) amplitudes (Fig. A1 C), and higher input resistance compared with CA1 neurons (Fig. A1 D). Although CA2 neurons showed higher input resistance than CA1 neurons, this difference was not significant when compared with FC neurons. In addition to putative excitatory neurons, we identified a population of interneurons within the FC based on their distinct electrophysiological properties (Fig. A2). These cells were identified based on their fast-spiking firing patterns (Fig. A2 A) and unique electrophysiological signatures (Fig. A2 B–C), including depolarized resting membrane potential (RMP; Fig. A2 E), hyperpolarized action potential threshold (Fig. A2 H), prominent afterhyperpolarizations (AHPs; Fig. A2 I), and increased input resistance and sag (Fig. A2 J-K).

**Figure 1.**
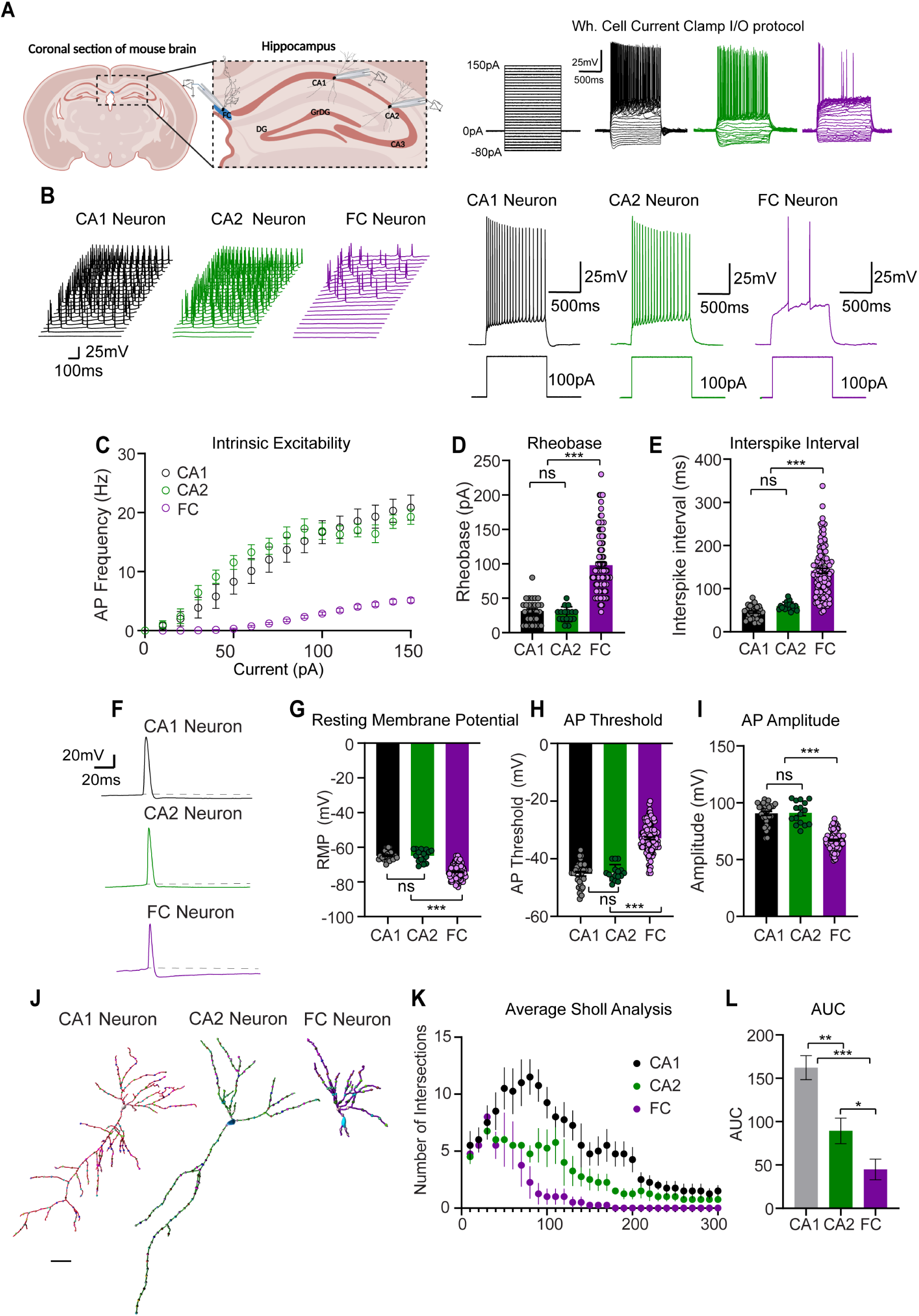
FC neurons exhibit reduced intrinsic excitability and distinct dendritic architecture compared with CA1 and CA2 neurons. (A) Schematic representation of the hippocampal slice showing the relative positions of CA1, CA2, and FC where whole-cell recordings were obtained and scheme of the current clamp protocol showing representative traces of CA1, CA2 and FC after progressive current injections from −80pA to 150pA. (B) Representative voltage traces from CA1, CA2, and FC neurons in response to current injection steps illustrating markedly lower firing in FC neurons. (C) Input–output relationship showing the number of action potentials (APs) as a function of injected current amplitude. (D–E) Quantification of rheobase (D; CA1 PNs= 31.3 ± 3 pA; CA2 PNs= 26.9 ± 2.7 pA; FC= 97.9 ± 4.1 pA) and interspike interval (E; CA1 PNs= 45.8 ± 2.7 ms; CA2 PNs= 59.3 ± 2.3 ms; FC= 141.2 ± 6 ms) demonstrating that FC neurons require stronger depolarizing currents to reach threshold and fire at lower frequencies. (F–I) Representative traces of single action potentials and comparison of passive and active membrane properties across regions: (G) resting membrane potential (RMP; CA1 PNs= −64.9 ± 0.5 mV; CA2 PNs= −64.9 ± 3.5 mV; FC= −74.1 ± 0.4 mV), (H) AP threshold (CA1 PNs= 45.2 ± 0.8 mV; CA2 PNs= −44.8 ± 0.7 mV; FC= −32.9 ± 0.4 mV) and (I) AP amplitude (CA1 PNs= 90.6 ± 1.7 mV; CA2 PNs= 91.1 ± 2.4 mV; FC= 67 ± 0.7 mV). (J) Representative neuron Imaris-tracing models used for Sholl analysis. (K) Sholl analysis of dendritic complexity generated in Imaris, showing reduced dendritic branching across radial distances in FC neurons compared with CA1 and CA2 pyramidal neurons. (L) Area under the curve (AUC; CA1 PNs= 162.3 ± 13.8 μm^2^; CA2 PNs= 89.4 ±14.6 μm^2^; FC= 44.9 ± 11 μm^2^) integrating dendritic intersections across distance for each cell. Data are presented as mean ± SEM; statistical comparisons were performed using one-way ANOVA followed by post hoc multiple comparisons (*p < 0.05, ** p < 0.001, \*\*\**p* < 0.0001). CA1 (n=30 cells/N=9 animals); CA2 (n=15/N 5); FC (n=117/N= 35).

To determine whether intrinsic physiological differences were associated with structural specialization, we performed reconstructions of biocytin-filled neurons from CA1, CA2, and the FC (Fig. 1 J). Quantitative morphological analysis revealed pronounced differences in dendritic architecture across regions. Sholl analysis demonstrated reduced dendritic complexity in FC neurons compared with both CA1 and CA2 neurons, particularly at distal radii (Fig. 1 K-L). This reduction was captured by a significantly smaller area under the Sholl curve (AUC) in FC neurons relative to CA1 and CA2. Together, these results indicate that FC neurons are characterized by reduced intrinsic excitability and simplified dendritic architecture relative to CA1 and CA2 neurons.

### FC neurons display heterogeneous intrinsic properties scaled with increased synaptic input

PCA of intrinsic excitability values (n. of APs) and input–output analysis revealed broad variability in FC firing behavior (Fig 2 A-C). To investigate the ionic mechanisms underlying this heterogeneity, we stratified FC neurons into low-excitability and high-excitability groups and compared their intrinsic properties. Low-excitability neurons (L), which comprised 60.7% of recorded FC cells (71/117 cells), displayed regular spiking with strongly reduced excitability. Compared with L, High-excitability (H) neurons exhibited higher firing frequencies in response to depolarizing current injections (Fig. 2 D–E), lower rheobase values (Fig. 2 F), and shorter interspike intervals (Fig. 2 G), indicating enhanced excitability. Other properties, including resting membrane potential and AP threshold, did not differ significantly between groups (Fig. 2 I–J), whereas high excitability FC neurons displayed smaller mAHPs (Fig. 2 L), higher input resistance (Fig. 2 N), and reduced sag amplitudes (Fig. 2 O).

**Figure 2.**
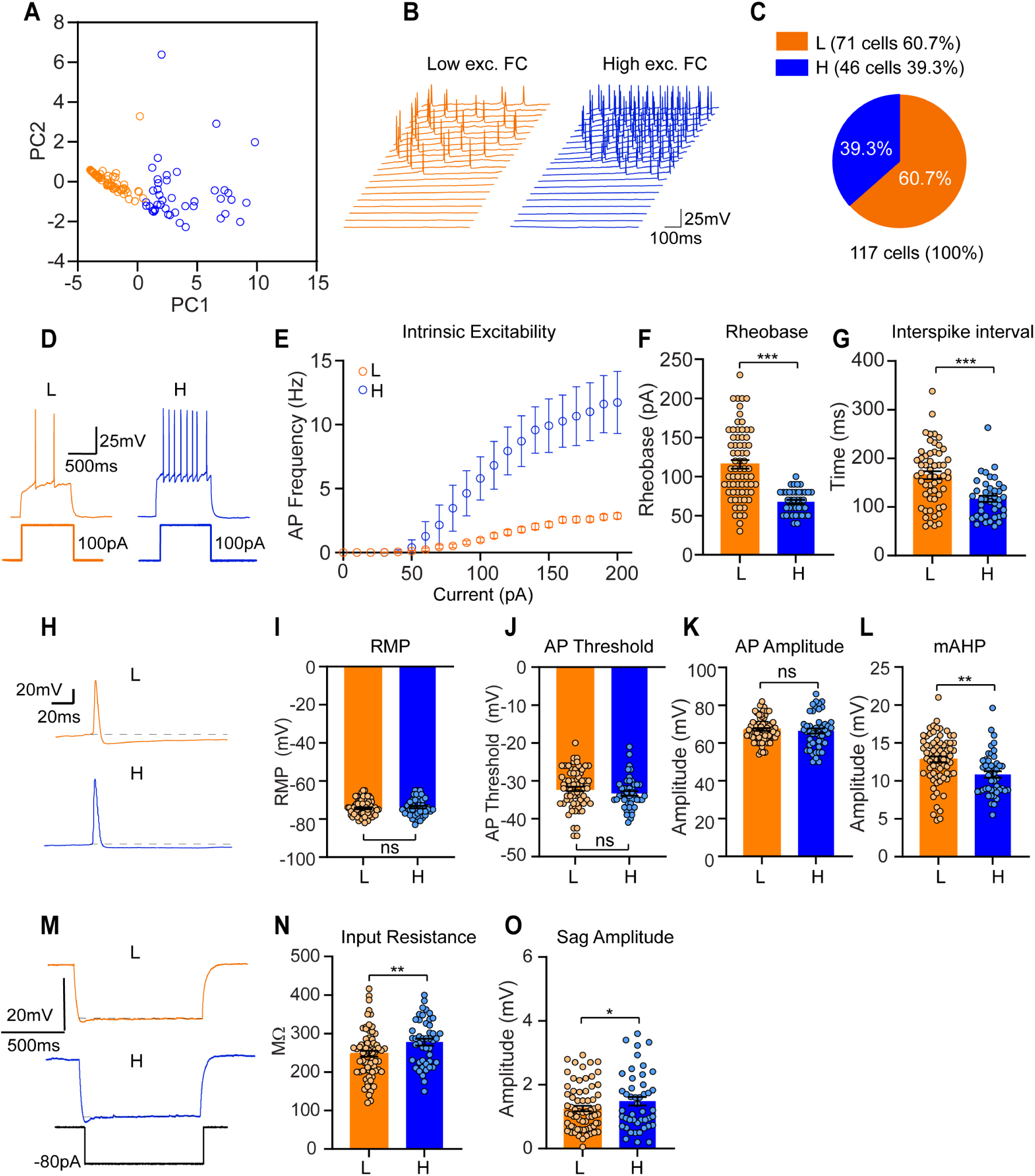
Continuum of intrinsic electrophysiological properties across FC neurons. (A) Principal component analysis (PCA) of intrinsic excitability parameters derived from input– output (I–O) relationships (number of action potentials across current injections) illustrating variability across the FC neuronal population. Neurons are color-coded according to relative excitability to visualize the distribution along the principal axis. (B) Distribution of key firing parameters across FC neurons demonstrating broad variability in intrinsic excitability. (C) Proportion of neurons represented across the range of excitability values within the FC population. (D–E) Representative voltage traces (D) and input–output relationships (E) from FC neurons spanning the range of intrinsic excitability, illustrating differences in firing responses during depolarizing current injections. (F–G) Quantification of rheobase (F; L= 115.7 ± 5.4 pA; H= 67.7 ± 2.4 pA) and interspike interval (G; L= 165.7 ± 8.1 ms; H= 116.8 ± 6.3 ms) across neurons, showing variability in intrinsic firing properties. (H–J) Representative single action potential traces (H) and comparison of intrinsic membrane properties, including RMP (I; L= −74.3 ± 0.5 mV; H= −73.7 ± 0.7 mV), AP threshold (J; L= −32.6 ± 0.3 mV; H= −33 ± 0.6 mV), AP amplitude (K; L= 67.2 ± 0.7 mV; H= 66.4 ± 1.4 mV), and medium afterhyperpolarization (mAHP; L; L= 12.84 ± 0.4 mV; H= 10.8 ± 0.4 mV). (M–O) Representative traces and quantification of input resistance (N; L= 247.5 ± 7.4 MΩ; H= 277.7 ± 8.7 MΩ) and hyperpolarization-induced sag (O; L= 1.25 ± 0.1 mV; H= 1.5 ± 0.1 mV), illustrating variability in passive and subthreshold membrane properties across FC neurons. For visualization, neurons at the lower and higher ends of the excitability spectrum (L and H) are indicated without implying discrete cellular subtypes. Data are presented as mean ± SEM; statistical comparisons were performed using unpaired two-tailed test (*p < 0.05, ** p < 0.001, \*\*\**p* < 0.0001). L (n=71 cells/N=30 animals); H (n=46/ N=20).

To assess whether differences in intrinsic excitability are accompanied by distinct synaptic input, we recorded spontaneous excitatory synaptic activity in FC and CA1 neurons (Fig. A3). FC neurons exhibited a significantly higher frequency of spontaneous excitatory postsynaptic currents (sEPSCs) compared with CA1 neurons (Fig. A3 A-C), indicating increased excitatory synaptic drive. Within the FC population, sEPSC frequency varied across neurons and was inversely related to intrinsic excitability, with lower-excitability neurons displaying higher event frequencies, whereas higher-excitability neurons exhibited lower excitatory synaptic input. Despite this variability, sEPSC frequency remained elevated in FC neurons overall compared with CA1. A small but significant difference in sEPSC decay time was also observed between CA1 and FC (Fig. A3 F). In contrast, sEPSC amplitude did not differ significantly across conditions (Fig. A3 E), suggesting that postsynaptic response properties are largely preserved. These findings indicate that excitatory synaptic input is enhanced in FC neurons and is differentially distributed across the population.

To further examine this relationship without imposing discrete classifications, we performed additional principal component analysis (PCA) incorporating a combined set of intrinsic parameters and synaptic properties across the FC population (Fig. A4 A-B). PCA of intrinsic features (Fig. A4 A) revealed a broad distribution of neurons, consistent with the variability observed in all firing behavior parameters, while PCA of synaptic parameters (Fig. A4 B) similarly showed broad variability. Importantly, comparison of these feature spaces revealed a structured continuum in which intrinsic excitability and synaptic drive were inversely related. Neurons with lower intrinsic excitability tended to exhibit higher spontaneous synaptic activity, whereas more excitable neurons received comparatively weaker input. Together, these findings indicate that FC neurons as a whole span a continuum defined by coordinated variation in intrinsic and synaptic properties.

These findings demonstrate that FC neurons exhibit substantial variability in intrinsic electrophysiological properties across the population. Furthermore, FC neurons receive increased excitatory synaptic input, which varies across neurons and may be coordinated with intrinsic excitability.

### Kv2-mediated conductances regulate intrinsic excitability in FC neurons

Given the distinctive electrophysiological features of FC neurons, including hyperpolarized resting membrane potentials, prominent afterhyperpolarizations, and markedly reduced excitability, we hypothesized that elevated potassium channel activity contributes to these properties. To test this, we recorded outward K⁺ currents using a voltage-clamp protocol (28–31). Kv-mediated currents varied substantially across FC neurons, paralleling the broad range of intrinsic excitability observed in current-clamp recordings (Fig. 3 A-B). To directly assess the relationship between potassium conductance and firing behavior, we performed correlation analysis between delayed rectifier K⁺ current amplitude and firing output. This analysis revealed a significant negative correlation (Fig. 3 C), indicating that neurons with larger Kv-mediated currents exhibit reduced excitability. These findings support a model in which graded differences in potassium conductance contribute to the heterogeneity of intrinsic firing properties across FC neurons.

**Figure 3.**
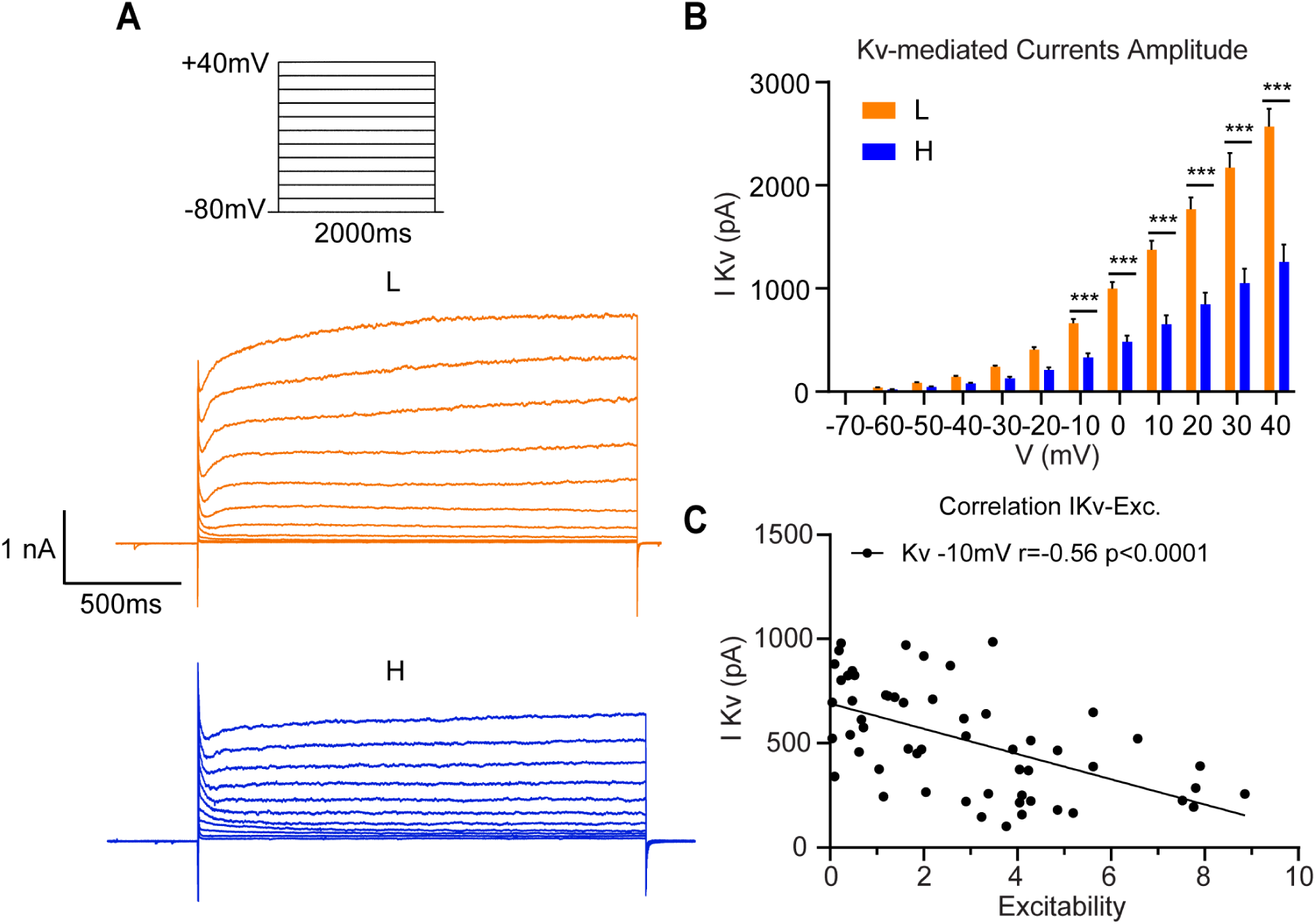
Variability of Kv-mediated currents across FC neurons and their relationship to intrinsic excitability. (A) Representative voltage-clamp recordings of outward potassium currents from FC neurons illustrating variability across the population. Neurons at the lower and higher ends of the excitability range are shown for comparison. (B) Quantification of Kv-mediated current amplitudes across FC neurons, demonstrating broad variability in delayed rectifier conductances. (C) Correlation between delayed rectifier K⁺ current amplitude and intrinsic firing properties (number of AP average), showing an inverse relationship between potassium conductance (at −10mV) and neuronal excitability. Unpaired Student t-test was used to compare current amplitudes between groups at each voltage step (***p<0.0001). L (n=32 cells/ N=13 animals); H (n=32/N=11). Summary values are outlined in Table A1.

Transcriptomic studies have previously identified high expression of AMIGO2 in the FC (32), a cell adhesion molecule known to interact with Kv2 (KCNB) channels and regulate their clustering and surface trafficking (28, 33). Kv2 channels are key determinants of delayed rectifier K⁺ currents and play a central role in action potential repolarization and firing rate modulation (29, 34, 35), suggesting a potential mechanism underlying the reduced excitability of FC neurons. To examine Kv2.1 channel distribution, we performed immunohistochemical labeling followed by ROI-based quantitative analysis. Kv2.1 immunoreactivity exhibited a punctate somatic pattern consistent with clustered surface expression in both CA1 and FC neurons (Fig. A5 A). Quantitative analysis revealed that FC neurons displayed significantly greater integrated fluorescence density and a larger Kv2.1-positive somatic membrane area compared with CA1 neurons (Fig. A5 C-E). In addition, the distribution of Kv2.1 signal in FC neurons was right-skewed, with a subset of neurons exhibiting particularly high values (Fig. A5 C, E), consistent with heterogeneous organization of Kv2-enriched membrane domains across the FC population.

To functionally assess the contribution of Kv2 channels to intrinsic excitability, we applied the Kv2-selective blocker Guangxitoxin-1E (GxTx). GxTx significantly reduced outward K⁺ currents across FC neurons (Fig. 4 C-F). To isolate the Kv2-dependent component, we performed point-by-point subtraction of averaged control and drug traces, yielding a GxTx-sensitive current with sustained activation kinetics characteristic of delayed rectifier conductances (Fig. A6 A). Quantification of the GxTx-sensitive current revealed substantial variability across neurons, consistent with the heterogeneity observed in baseline Kv-mediated currents (Fig. A6 B–C). Importantly, the amplitude of the GxTx-sensitive component was associated with intrinsic excitability, indicating that Kv2-mediated conductance contributes to the observed variability in firing behavior. Consistent with these voltage-clamp findings, GxTx application increased intrinsic excitability and reduced rheobase across FC neurons (Fig. 5 B–D). These effects were accompanied by increased input resistance (Fig. 5 J), reduced medium afterhyperpolarization amplitude (Fig. 5 I), and shifts in resting membrane potential and action potential threshold (Fig. 5 F–G). Together, these results demonstrate that Kv2 channels exert a strong suppressive influence on intrinsic excitability in FC neurons and contribute to the heterogeneity of their firing properties.

**Figure 4.**
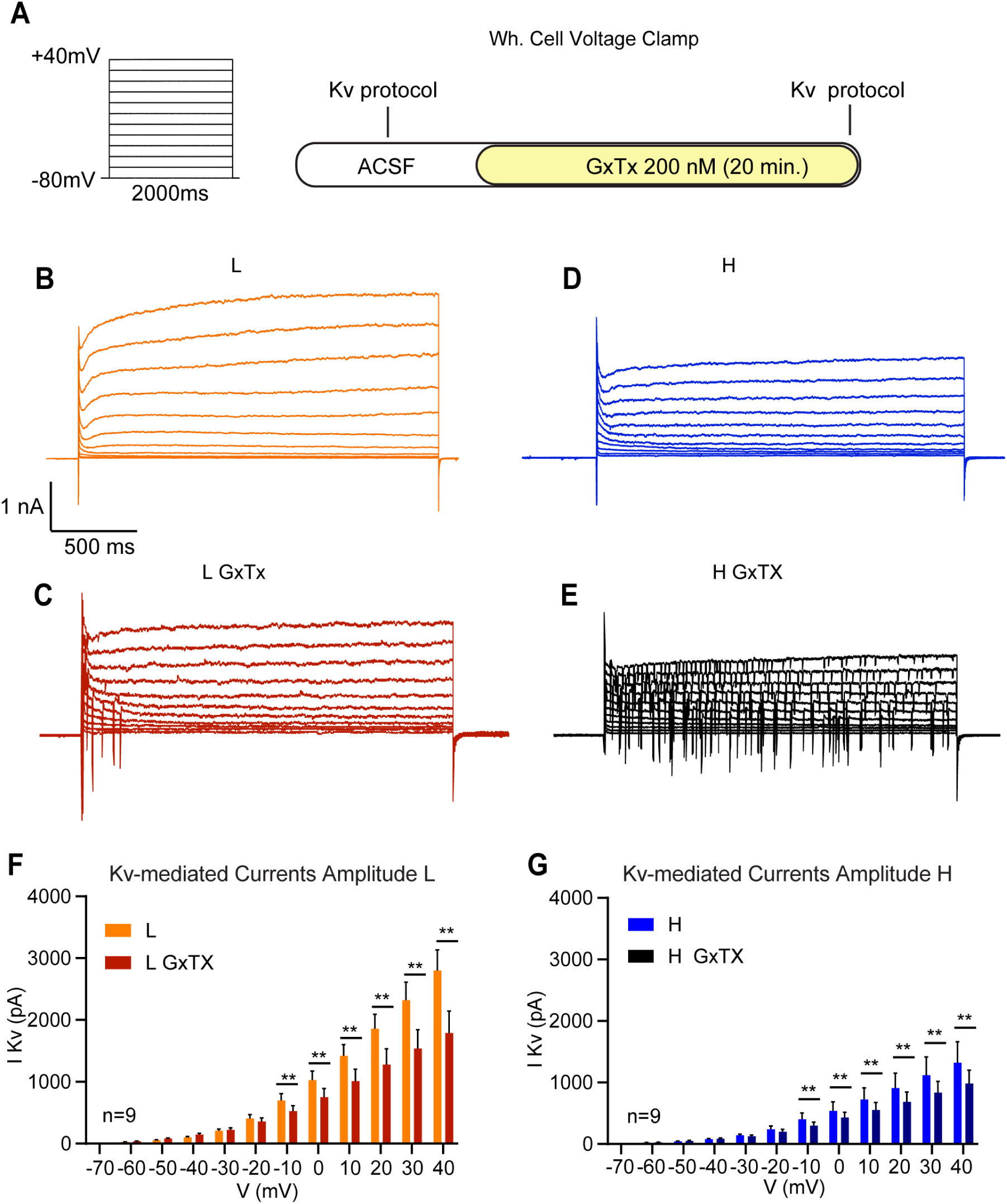
GxTx-sensitive potassium currents contribute to intrinsic excitability in FC neurons. (A) Voltage-clamp protocol used to evoke delayed rectifier potassium currents in FC neurons. Cells were held at −80 mV and stepped to depolarized potentials up to +40 mV (2000 ms duration) under control conditions (ACSF) and following bath application of Guangxitoxin-1E (GxTx, 200 nM). (B, D) Representative families of outward potassium currents recorded from FC neurons at the lower (L; B) and higher (H; D) ends of the excitability range under control conditions. (C, E) Representative current traces from the same neurons shown in (B, D) following GxTx application, demonstrating a reduction in outward current amplitude. (F, G) Quantification of Kv-mediated current amplitudes as a function of membrane potential for neurons at the lower (F) and higher (G) ends of the excitability range, before and after GxTx application. GxTx significantly reduced outward current amplitudes across depolarized voltage steps, with variability across neurons. A paired Wilcoxon signed-rank test was used to compare current amplitudes at each voltage step before and after GxTx application (**p < 0.01). L non-treated (n=9 cells/N=8 animals); L GxTx (n=9/N=7); H non-treated (n=9/N=8); H GxTx (n=9/N=7). Summary values are outlined in Table A2.

**Figure 5.**
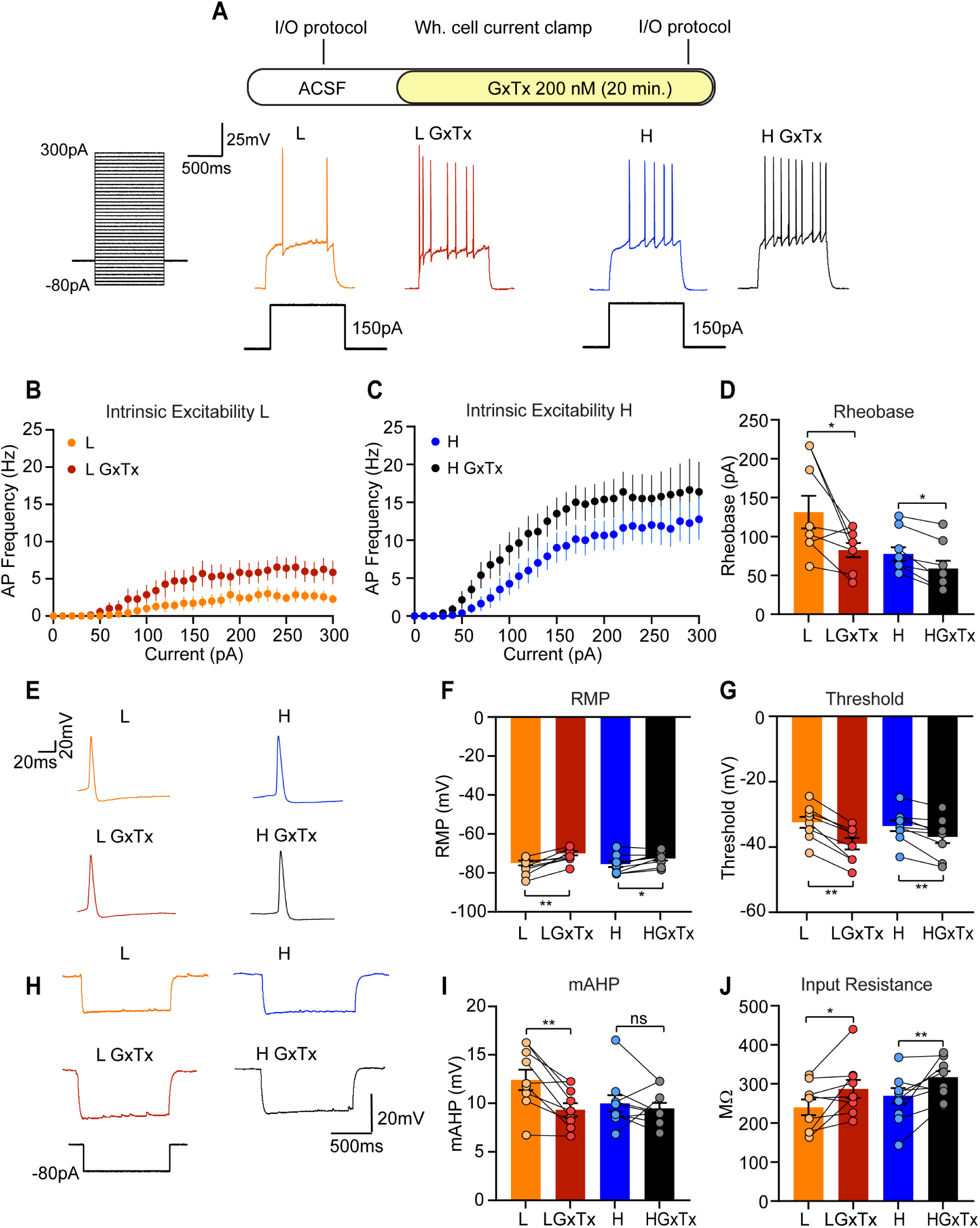
Kv2 channel blockade increases intrinsic excitability in FC neurons. (A) Experimental schematic and representative whole-cell current-clamp recordings from FC neurons during input–output (I/O) protocols in control conditions (ACSF) and following bath application of Guangxitoxin-1E (GxTx, 200 nM, 20 min). Example voltage responses from neurons at the lower (L) and higher (H) ends of the excitability range are shown before and after GxTx application. (B–C) Input–output relationships showing action potential (AP) firing frequency as a function of injected current in control conditions and after GxTx application for neurons with lower (B) and higher (C) intrinsic excitability. GxTx increases firing frequency across current injections. (D) Quantification of rheobase demonstrating a reduction following GxTx application across neurons (L= 131.3 ± 21 pA; L GxTx= 82.5 ± 9.2 pA; H= 77.5 ± 8.8 pA; H GxTx= 58.7 ± 10 pA). (E) Representative single action potential traces from FC neurons before and after GxTx application. (F–G) Summary of RMP (F; (L= −74.9 ±1.2 mV; L GxTx= −69.9 ± 1 mV; H= −75.5 ± 1.4 mV; H GxTx= −72.5 ± 1 mV) and AP threshold (G; (L= −32.4 ±1.7 mV; L GxTx= −39 ± 1.7 mV; H= −33.5 ± 1.6 mV; H GxTx= −36.9 ± 1.9 mV), showing depolarization of RMP and hyperpolarization of threshold following Kv2 channel blockade. (H) Representative voltage responses to hyperpolarizing current steps illustrating changes in membrane properties after GxTx application. (I–J) Quantification of mAHP (I; L= 12.4 ±1 mV; L GxTx= 9.3 ± 0.7 mV; H= 10 ± 0.8 mV; H GxTx= 9.4 ± 0.5 mV) and input resistance (J; L= 239.6 ± 20 MΩ; L GxTx= 286.7 ± 23 MΩ; H= 269.2 ± 19 MΩ; H GxTx= 316.7 ± 15.4 MΩ), showing reduced mAHP amplitude and increased input resistance following GxTx application. Data are presented as mean ± SEM; a paired Wilcoxon signed-rank test was used to compare statistically all parameters before and after GxTx application (* p<0.01; **p<0.001). L non-treated (n=9 cells/N=8 animals); L GxTx (n=9/N=8); H non-treated (n=9/N=8); H GxTx (n=9/N=8).

### Kv7-mediated conductances contribute to variability in intrinsic excitability across FC neurons

The hyperpolarized resting membrane potential of FC neurons suggests that potassium channels active at subthreshold potentials may also contribute to their intrinsic excitability. Among these, Kv7 (KCNQ) channels mediate the M-current, a non-inactivating subthreshold conductance that stabilizes membrane potential and suppresses repetitive firing. Kv7.2 and Kv7.3 subunits are widely expressed in the hippocampus and exert strong inhibitory control over neuronal excitability (36–38).

To assess the contribution of Kv7 channels in FC neurons, we recorded M-currents under voltage-clamp conditions. Kv7-mediated currents exhibited substantial variability across FC neurons, paralleling the heterogeneity observed in intrinsic excitability (Fig. 6 B–C, F). Notably, neurons with higher intrinsic excitability tended to display smaller M-current amplitudes, suggesting a reduced contribution of subthreshold potassium conductance in these cells. Importantly, Pearson correlation analysis revealed a significant negative correlation between Kv7-mediated current amplitude and firing output (Fig. 6 I).

**Figure 6.**
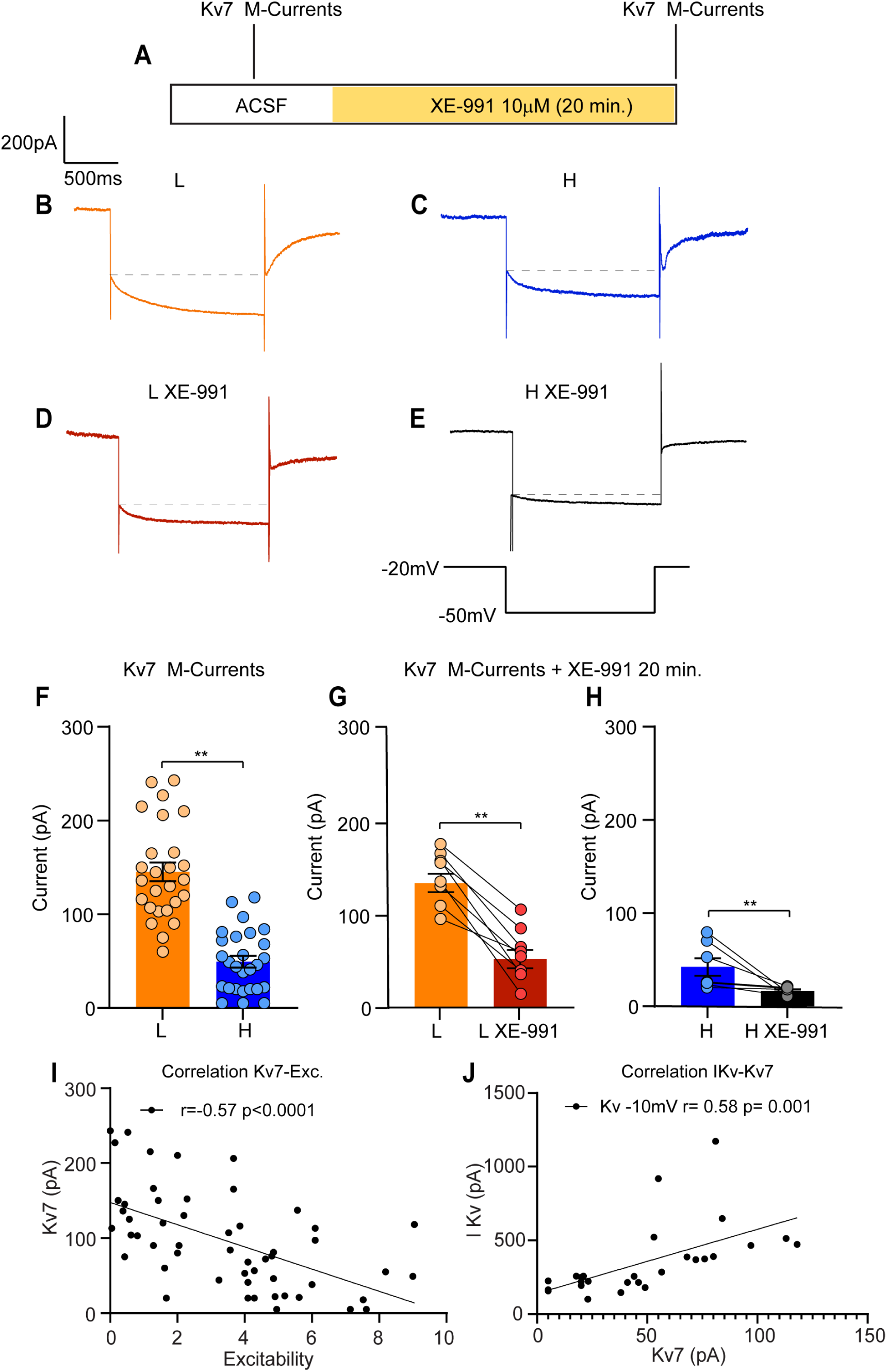
Kv7-mediated M-currents vary across FC neurons and are associated with intrinsic excitability and Kv2 conductance. (A) Experimental protocol for recording M-currents, showing bath application of the Kv7 channel blocker XE-991 (10 μM, 20 min). (B–C) Representative M-current traces from FC neurons spanning the range of intrinsic excitability, illustrating larger currents in neurons with lower excitability (L) and smaller currents in more excitable neurons (H). (D–E) Representative traces recorded after XE-991 application, demonstrating reduction of M-currents in both neurons. (F) Quantification of M-current amplitude across FC neurons, showing variability and reduced currents in more excitable neurons (L= 145 ± 10 pA; H = 49.3 ± 6.2 pA). (G) Comparison of M-current amplitude before and after XE-991 application in neurons at the lower end of the excitability range, demonstrating significant current reduction (L= 135 ± 10 pA; L XE-991 = 52 ± 10 pA) (H) Paired comparison of M-current amplitude before and after XE-991 in neurons at the higher end of the excitability range, showing effective blockade (H= 43 ± 9 pA; H XE-991 = 16 ± 1.5 pA). (I) Kv7-mediated current amplitude is inversely correlated with intrinsic excitability, indicating that reduced M-current is associated with increased firing output across FC neurons. (J) Correlation between Kv7-mediated M-current amplitude and rectifying Kv-mediated currents amplitude across FC neurons, revealing a significant positive relationship (Pearson correlation). Kv7-mediated M-current amplitude positively correlates with Kv2-mediated current amplitude, suggesting coordinated regulation of subthreshold and delayed rectifier potassium conductances. Data are presented as mean ± SEM; a paired Wilcoxon signed-rank test was used to compare statistically all parameters (**p<0.001). L (n=27 cells/N=12 animals); H (n=27/N=12); L before XE (n=8/N=5); L after XE (n=8/N=5); H before XE (n=7/N=5); H after XE (n=7/N=5). For panel I, n=53/N=15; for panel J, n=27/N=6.

To further examine the functional contribution of Kv7 channels, we applied the selective Kv7 blocker XE-991. XE-991 produced a robust reduction in M-currents across FC neurons (Fig. 6 D–E, G–H), confirming the identity of the recorded current. The magnitude of the blockade varied across neurons, consistent with differences in baseline M-current amplitude. Interestingly, Kv7-mediated currents were positively correlated with delayed rectifier potassium currents across FC neurons (Fig. 6 J), suggesting coordinated regulation of subthreshold (Kv7-mediated) and delayed rectifier (Kv2-mediated) potassium conductances. Together, these findings indicate that variability in Kv7 channel function contributes to the heterogeneity of intrinsic excitability in FC neurons and acts in concert with Kv2-mediated conductances to shape firing behavior.

## Discussion

Our results identify the fasciola cinereum (FC) as a distinct hippocampal neuronal population characterized by reduced intrinsic excitability and unique structural features. FC neurons differ from neighboring CA1 and CA2 neurons in both electrophysiological and morphological properties, supporting the view that the FC is not a dorsal extension of these regions but a separate circuit element with specialized cellular characteristics.

Although FC shares many molecular markers with CA2 such as PCP4 and AMIGO2, our findings demonstrate that its physiological profile is fundamentally distinct. CA2 pyramidal neurons exhibit depolarized resting membrane potentials, minimal spike-frequency adaptation, and high firing rates associated with rapid encoding functions (39–41). In contrast, FC neurons exhibited pronounced spike-frequency adaptation, large afterhyperpolarizations, and hyperpolarized resting membrane potentials, indicating substantially reduced intrinsic excitability. Importantly, input resistance in FC neurons was not lower than in CA2 or CA1 neurons, suggesting that their hypoexcitability stems from enhanced active potassium conductances rather than passive membrane differences. Together, these comparisons underscore that FC and CA2 are electrophysiologically distinct. In addition to their unique electrophysiological properties, FC neurons exhibited differences in dendritic organization compared to neighboring hippocampal pyramidal neurons, with reduced dendritic complexity at distal radii. These features may influence synaptic integration and contribute to the functional profile of FC neurons. However, these observations should be interpreted with caution, as reconstructions from single acute slices may underestimate dendritic extent due to truncation, and differences in slicing orientation between regions may affect dendritic preservation. Together these results suggest that the FC represents a functionally and anatomically distinct hippocampal subregion.

FC neurons exhibit substantial heterogeneity in intrinsic electrophysiological properties. In our analysis we segregated FC neurons into low and high excitability groups, although we acknowledge that this grouping does not mean that these are distinct subtypes, and our data may still be significantly underpowered to resolve all cell types in FC. This variability in excitability is closely associated with differences in potassium channel–mediated conductances. In particular, delayed rectifier currents mediated by Kv2 channels varied across FC neurons and were inversely related to intrinsic excitability. Kv2 channels are known to localize to somatic membrane domains where they exert strong control over spike repolarization and firing dynamics (42, 43). Consistent with this, our immunohistochemical analysis indicates that FC neurons exhibit a greater Kv2.1-positive membrane area and increased integrated Kv2 signal compared to CA1 neurons, suggesting an expansion of functional Kv2.1-enriched membrane domains at the soma (44). Although this analysis does not resolve the number or size of individual channel assemblies, increased integrated density over a larger somatic area is expected to enhance the effective contribution of Kv2-mediated delayed rectifier currents. This organization provides a plausible structural correlate for the elevated rheobase, prolonged interspike intervals, and pronounced afterhyperpolarizations observed electrophysiologically in FC neurons.

Pharmacological blockade of Kv2 channels with GxTx resulted in consistent depolarization of the resting membrane potential, decreased rheobase, and increased input resistance, collectively indicating a reduction in Kv2-mediated K⁺ conductance. The observed increase in input resistance after GxTx supports a genuine decrease in resting K⁺ conductance, reinforcing the notion that Kv2 channels provide tonic outward current that constrains excitability (31). Kv2 channels have been shown to regulate neuronal excitability in dissociated neuronal preparations (30). Here, we extend these findings to FC neurons in acute hippocampal slices, demonstrating their contribution to intrinsic firing properties in this understudied subregion. Although voltage-clamp protocols involving prolonged depolarization may permit Ca²⁺ influx and potential modulation of ion channels, the pharmacological sensitivity of the recorded currents supports a dominant contribution of Kv-mediated conductances. These findings underscore the functional importance of Kv2 conductances in shaping intrinsic excitability in FC neurons and suggest that they may represent potential targets in conditions characterized by altered neuronal excitability, such as epilepsy (45, 46).

In addition to Kv2, we identified a substantial M-current in FC neurons, mediated by Kv7 channels. These channels provide tonic inhibition and support spike-frequency adaptation, stabilizing the resting membrane potential and suppressing repetitive firing (37, 47), both in hippocampal and cortical neurons (38, 48). Kv7-mediated currents also varied across FC neurons and were associated with intrinsic excitability, with smaller M-currents observed in more excitable neurons. Given the direct association of Kv7 mutations with epileptic encephalopathies and the clinical efficacy of Kv7 channel openers such as retigabine, these channels may serve as critical targets for modulating FC function under pathological conditions (49–51). The positive correlation between Kv2 and Kv7 function in FC neurons suggests a coordinated expression or functional coupling between these two potassium channel families, which together shape the excitability landscape of FC neurons.

Recent *in vivo* studies have reported divergent findings regarding FC neuron firing. Recordings in freely behaving animals indicate sparse and delayed firing consistent with reduced excitability (12), whereas juxtacellular recordings under anesthesia report firing properties more similar to CA1 (11). These discrepancies may reflect differences in network state. Under anesthesia, reduced synaptic drive may mask intrinsic differences in excitability, whereas awake behavioral conditions engage stronger network activity. Notably, we found that FC neurons receive substantial excitatory synaptic input, with significantly increased sEPSC frequencies compared to CA1, yet they do not exhibit correspondingly high firing rates either *in vivo* from previous studies or *ex vivo* from our results. Within the FC population, synaptic input was inversely related to intrinsic excitability, with lower-excitability neurons receiving higher-frequency excitatory input. Our additional PCA analyses further support this relationship by showing that FC neurons occupy a continuum of electrophysiological states, structured along an axis defined by intrinsic excitability and synaptic drive, suggesting coordinated variation in intrinsic and synaptic properties. Given that intrinsic excitability in FC neurons is strongly governed by potassium conductances, particularly Kv2- and Kv7-mediated currents, these findings suggest that variability in ion channel composition defines the position of individual neurons along this functional axis. Neurons with stronger potassium conductances exhibit reduced intrinsic excitability and are associated with increased synaptic input, whereas more excitable neurons receive comparatively weaker synaptic drive. This relationship suggests a potential compensatory organization in which increased synaptic drive is balanced by stronger intrinsic constraints on excitability. This interplay between synaptic input and intrinsic conductances provides a potential explanation for discrepancies across *in vivo* studies, where differences in network state may differentially engage these intrinsic constraints. These findings highlight how intrinsic membrane mechanisms can impose strong constraints on neuronal activity downstream of circuit connectivity strength. The Kv2- and Kv7-mediated conductances identified here are well positioned to limit spike output even in the presence of increased synaptic drive. An understanding of these fundamental properties will be critical for future studies linking circuits, activity, and behavioral output in the FC.

Recent work has identified the fasciola cinereum as a critical node in seizure initiation and propagation, with both human intracranial recordings and animal models demonstrating a causal role for FC activity in epilepsy (18). While these studies establish the functional importance of the FC at the circuit level, the cellular mechanisms underlying this role have remained unclear. Our findings provide a potential biophysical framework linking intrinsic membrane properties to this function. The inverse relationship we observe between intrinsic excitability and synaptic drive suggests a coordinated organization in which increased input is balanced by intrinsic suppression of firing. In this context, perturbations of potassium channel function or synaptic balance could shift FC neurons toward hyperexcitability, potentially facilitating seizure initiation or propagation. Although direct circuit-level validation will be required, these results identify intrinsic and synaptic properties of FC neurons which may contribute to its previously identified role in epilepsy.

Altogether, our results indicate that the fasciola cinereum is not merely a transitional zone but a distinct hippocampal subregion with specialized intrinsic properties. The combination of reduced intrinsic excitability, compact dendritic architecture, and prominent potassium channel– mediated conductances suggests that FC neurons operate under strong intrinsic control and are organized along a functional axis in which intrinsic excitability and synaptic input are coordinated. These features may contribute to shaping how synaptic inputs are integrated within the hippocampal circuit and provide a framework for understanding how intrinsic excitability influences seizure propagation and network dynamics. Future studies will be required to determine how these intrinsic properties interact with synaptic and circuit-level activity to regulate FC function *in vivo*.

## Materials and Methods

### Mice

All experimental procedures were approved by the Institutional Animal Care and Use Committee (IACUC) at Vanderbilt University and were conducted in accordance with the National Institutes of Health guidelines for the care and use of laboratory animals. C57BL/6J mice of either sex (postnatal day 21–60) were used for all experiments. Animals were housed under standard conditions (12 h light/dark cycle, food and water ad libitum) and monitored daily for health and well-being. Every effort was made to minimize animal suffering and the number of animals used.

### Hippocampal Slice Preparation

Coronal hippocampal slices (280 µm) were prepared from postnatal day (P) 21–60 from C57BL/6J mice of either sex. Mice were deeply anesthetized with isoflurane and euthanized by decapitation. Brains were rapidly removed and placed in ice-cold, oxygenated N-methyl-D-glucamine (NMDG)-based cutting solution containing (in mM): 92 NMDG, 1.2 NaH₂PO₄, 30 NaHCO₃, 20 HEPES, 25 glucose, 5 sodium ascorbate, 3 sodium pyruvate, 10 MgCl₂, and 0.5 CaCl₂ (pH adjusted to 7.3– 7.4, 300–310 mOsm). Coronal slices were cut using a vibratome (Leica VT-1200S) and transferred to a holding chamber containing oxygenated artificial cerebrospinal fluid (ACSF) at room temperature (24–26°C) for at least 1 h before recording. ACSF contained (in mM): 125 NaCl, 2.5 KCl, 0.8 NaH₂PO₄, 26 NaHCO₃, 3 CaCl₂, 2 MgCl₂, and 10 glucose, continuously bubbled with 95% O₂/5% CO₂.

To enable reliable targeting of CA2 pyramidal neurons, hippocampal slices were prepared following anatomical and electrophysiological criteria described by Wei and colleague (52), with adaptations to our recording procedures. The hippocampus was rapidly dissected and transferred to ice-cold cutting solution. Slices (300 µm) were prepared in the horizontal plane using a vibrating microtome, an orientation selected to preserve the anatomical and synaptic connections spanning CA2. After recovery in ACSF, slices were transferred to the recording chamber and visualized under IR-DIC optics. CA2 pyramidal neurons were identified using a combination of positional landmarks, cytoarchitectonic features, and intrinsic membrane properties.

Anatomically, CA2 neurons were targeted within the narrow sector of the stratum pyramidale situated near the dorsal CA1, adjacent to the end of the suprapyramidal blade of the dentate gyrus. Care was taken during slicing to preserve the CA3–CA2–CA1 boundary, which is highly sensitive to mechanical disruption. Neurons were selected only if they were located within this CA2-enriched zone and displayed characteristic CA2 somatic morphology (large pyramidal soma, sparse apical tufting relative to CA1).

Electrophysiological features previously established as hallmarks of CA2 neurons were used to further confirm identity, including relatively high input resistance, intermediate firing frequency and pronounced hyperpolarization-activated sag.

Only neurons fulfilling all anatomical and physiological criteria were classified as CA2 and included in the analysis.

### Electrophysiology

Recordings were performed at room temperature (24-26° C) in a recording chamber continuously perfused with oxygenated ACSF.

Neurons were recorded from the fasciola cinereum (FC), a small hippocampal subregion located at the dorsomedial edge of the hippocampal formation. The FC was identified in acute coronal slices based on its anatomical position using the Allen Mouse Brain Atlas as a reference (Bregma 1.67 mm to 2.03 mm), together with previously published descriptions (11, 12, 16, 18). Specifically, recordings were restricted to the region adjacent to the third ventricle and medial to the CA1 pyramidal layer, consistent with established FC boundaries.

To minimize potential contamination from neighboring regions, cells located within or near the CA2 pyramidal layer were systematically excluded. Only neurons within the visually defined FC region were included for analysis. The location of recorded neurons was verified based on anatomical landmarks and, when available, biocytin-based morphological reconstruction.

Whole-cell patch-clamp recordings were obtained using borosilicate glass pipettes (5–7 MΩ) filled with an internal solution containing (in mM): 120 K-gluconate, 20 KCl, 10 HEPES, 0 EGTA, 2 MgCl₂·6H₂O, 2 Na₂ATP, and 0.1 spermine (pH adjusted to 7.3 with KOH, ∼290 mOsm). Internal solution contained biocytin for *post-hoc* characterization. Following the formation of a high-resistance seal (>1 GΩ), gentle suction was applied to achieve the whole-cell configuration. Recordings were accepted only if access resistance (Ra) was ≤25 MΩ, leak currents were minimal, and the resting membrane potential was stable and hyperpolarized.

Cells were initially recorded in current-clamp mode to assess intrinsic membrane properties. Apparent input resistance was calculated using hyperpolarizing current steps (−80 pA, 800 ms). Series resistance was monitored throughout the experiment, and recordings were discarded if resistance changed by more than 10%. To assess excitability, action potentials (APs) were evoked using a series of incremental depolarizing current injections (10 pA steps, 1000 ms duration), and input–output curves were generated based on AP counts. Action potential threshold, amplitude, and afterhyperpolarization were extracted using automated routines based on first derivative thresholding.

Following current-clamp recordings, cells were returned to voltage-clamp mode to isolate and measure specific K⁺ currents. Kv-mediated delayed rectifier K⁺ currents were evoked using a series of 2000 ms voltage steps from −80 mV (or −70 mV) to +40 or +50 mV. To isolate the Kv2-dependent component of the delayed rectifier current, Guangxitoxin-1E (GxTx; 200 nM) was bath-applied following acquisition of stable baseline recordings. Identical voltage-step protocols were repeated at steady state (15–20 min after application). Currents were baseline-corrected prior to analysis. The GxTx-sensitive current was obtained by point-by-point subtraction of averaged control and drug traces (I_sensitive = I_control − I_GxTx). Quantification was performed at +20 mV, a voltage at which delayed rectifier currents were robustly activated and reached steady state under our recording conditions. Steady-state amplitudes were measured after 300 ms of each voltage step to minimize contributions from capacitive transients and fast inactivating components. Recordings were included only if series resistance remained stable (<10% change). Baseline recordings were obtained in the absence of pharmacological agents to enable within-cell comparisons following drug application.

M-currents were assessed using a standard deactivation protocol: cells were held at −15/20 mV and stepped to more hyperpolarized potentials (e.g., −50mV) as described by Lawrence et al. (2006). The relaxing component of the deactivating tail current was quantified as a measure of M-current amplitude.

### Principal component analysis (PCA) of intrinsic excitability

Principal component analysis (PCA) was applied to electrophysiological data derived from the input–output (I-O) current-clamp protocol in FC neurons (number of action potentials across depolarizing current steps). For each FC neuron, the number of action potentials generated at each depolarizing current step was used as a feature. Prior to PCA, all features were standardized by z-scoring. Two principal components were used for visualization which overall accounted for 90.96% of the variance (PC1: 67.61% variance, PC2: 23.35% variance) of the distribution of neurons in reduced feature space. The 70-200 pA range contributed most to PC1, while the 20-60 pA range contributed most to PC2. The 0 pA and 10 pA features did not contribute to either component. To facilitate comparison across the range of excitability profiles, k-means clustering (k = 2) was applied to the two principal components. This grouping was used solely as an analytical framework to contrast neurons at the lower and higher ends of the excitability spectrum and does not imply the existence of discrete neuronal subtypes.

Cluster assignments were visualized in the two-dimensional PCA space. Subsequent analyses of intrinsic membrane properties and firing behavior were performed using this excitability-based grouping to illustrate variability across the population.

### Principal component analysis of intrinsic and synaptic properties

To examine the relationship between intrinsic properties and synaptic input across FC neurons, we performed two PCAs on a combined set of electrophysiological parameters. Input resistance, rheobase, RMP, sag amplitude, AP amplitude, AP threshold, mAHP, and average I-O spike count were used as features to represent intrinsic electrophysiological properties (PC1: 29.05% variance, PC2: 20.97% variance). sEPSC frequency, inter episode interval length, amplitude, and decay time were used as features to represent synaptic properties (PC1: 41.23% variance, PC2: 26.99% variance). All features were z-score standardized and two principal components were used for visualization for each PCA. These analyses allowed identification of the dominant axes of variability across the FC population. No clustering algorithms were applied in this analysis. Instead, the distribution of neurons in principal component space was used to assess whether intrinsic and synaptic properties were organized into discrete groups or varied continuously across the population. For visualization, the same low and high excitability cells noted in the PCA of intrinsic excitability were also noted in these graphs.

### Pharmacology

To pharmacologically dissect channel-specific contributions to intrinsic excitability, the following compounds were bath-applied:

- XE991 (10 µM), a selective Kv7 (KCNQ) channel blocker (Tocris).
- Guangxitoxin-1E (GxTx, 200 nM), a potent and selective inhibitor of Kv2.1/2.2 channels (Alomone labs).

Drug effects were monitored continuously, and recordings were performed at steady-state (∼15– 20 min after drug application) to ensure complete superfusion into the slice.

### Data Acquisition, Analysis and Statistics

Signals were acquired using a Multiclamp 700B amplifier (Molecular Devices), digitized at 10– 20 kHz with a Digidata 1550B interface, and low-pass filtered at 2–5 kHz. Data were recorded and analyzed using pClamp software (Molecular Devices) and routines in the Easy Electrophysiology software. All data was processed for statistical analysis and figure presentation using Microsoft Excel and Prism GraphPad 10 software. Adobe Illustrator was used to build the final figures.

All data are reported as mean ± SEM. To compare intrinsic properties across CA1, CA2 and FC a one-way ANOVA was performed, followed by Tukey’s post hoc test for multiple comparisons. Normality of the data was assessed using the Shapiro-Wilk test prior to analysis. To compare FC G1 to G2 intrinsic properties, Kv2 and M-current amplitudes between neuronal groups, we used a two-tailed unpaired *t*-test when data passed the Shapiro–Wilk normality test; otherwise, a non-parametric Mann–Whitney U test was applied. For paired pharmacological experiments (e.g., before and after drug application within the same neuron), we applied for the Wilcoxon signed-rank test. Normality was assessed using the Shapiro-Wilk test, and statistical significance was set at p < 0.05.

### Tissue Collection, Fixation, and Sectioning

Mice were deeply anesthetized with isoflurane and transcardially perfused with 45 mL ice-cold 4% paraformaldehyde (PFA) pre-filtered through double-wrapped filter paper. Whole brains were collected, post-fixed for 24 h in 4% PFA at 4 °C, and then transferred to 1× PBS at 4 °C until further processing. Coronal sections were cut at 50 µm using a Leica VT1000S vibratome. Sections containing the fasciola cinereum (FC) were collected from anterior to posterior, yielding ∼10 slices per animal. Three interleaved free-floating series were collected in 1× PBS, immediately transferred to 1× TBS, and rinsed four times for 10 min before immunohistochemical processing.

### Immunohistochemistry for Kv2.1

After the initial TBS rinses, free-floating FC sections were incubated for 1 h at room temperature (RT) in TBS+ (1× TBS with 1% BSA and 0.05% Triton X-100) to block non-specific binding. Sections were then incubated for at least 17 h at RT on a gentle rocker in TBS+ containing one of the following primary antibodies: rabbit anti-Kv2.1 (Alomone Labs, Cat# APC-012; polyclonal; 1:2000).

Following primary incubation, sections were washed in 1× TBS (4 × 10 min) and incubated for 2 h at RT with CF®594 highly cross-adsorbed goat anti-rabbit IgG (H+L) secondary antibody (Biotium, Cat# 20113; 1:1000 in TBS+). Sections were then washed in 1× TBS (4 × 10 min), mounted onto Superfrost Plus charged slides, air-dried, and coverslipped with VECTASHIELD Antifade Mounting Medium with DAPI. Slides were sealed with nail polish, and no more than six sections were mounted per slide to maintain signal integrity. All staining experiments were performed with at least three biological replicates per group and at least three technical replicates per condition.

### Immunohistochemistry Quantification

Confocal images were acquired using a laser-scanning confocal microscope (Zeiss LSM 880) equipped with a 40× oil-immersion objective. To ensure comparability across samples, all images were collected using identical acquisition parameters, including laser power, detector gain, pinhole size, and exposure settings. Z-stacks were acquired at 0.5 µm intervals and processed uniformly using Fiji/ImageJ (NIH), minimizing bias in visualization and downstream quantification of signal distribution within the fasciola cinereum (FC) and adjacent hippocampal regions.

Quantification of Kv2.1 immunoreactivity was performed using ImageJ. For each section, three anatomically matched regions of interest (ROIs) were selected from the FC and neighboring CA1. Images were background-subtracted using the *Subtract Background* function (rolling ball radius: 100 pixels) to enhance contrast between Kv2.1-positive clusters and surrounding neuropil. A binary mask was then generated by applying a consistent intensity threshold to the Kv2 channel, selectively highlighting somatic and proximal dendritic Kv2.1-positive clusters. This mask was applied to the original image to isolate Kv2.1-positive signal. The *Analyze Particles* function was used to extract integrated fluorescence density and total Kv2.1-positive ROI area for each region. All analyses were conducted using identical thresholding and processing parameters across slices and experimental groups to ensure consistency.

### Biocytin filling and confocal microscopy

The morphology of the recorded neurons was revealed by biocytin staining. For this, biocytin (0.2– 0.4% ∼5.0 mM, Sigma-Aldrich) was added to the pipette solution, and cells were filled at least 20 min. Biocytin was revealed with streptavidin complex coupled to Alexa Fluor 488 (Thermo Fisher Scientific). Confocal z-stacks were acquired using a 20× or 40x oil immersion objective with a z-step of 0.5 µm on a laser-scanning confocal microscope (Zeiss LSM 880).

### Morphological reconstruction and analysis

Z-stack image data were imported into Imaris (version 10.2.0; Oxford Instruments) by converting Zeiss CZI files to Imaris IMS format using the Imaris File Converter (version 10.2.0). To reduce background noise and improve image quality prior to filament reconstruction, local background subtraction was performed in Imaris using a filter width of 53.1 µm.

Neuronal morphology was reconstructed using the AI Filament Tracer module in Imaris to detect and segment biocytin-labeled neuronal processes (Alexa Fluor 488). Soma detection was performed first to identify the neuronal cell body and define the origin of the filament tree. Automated filament detection was then carried out using the AutoPath (no loops, without spines) algorithm. To improve segmentation accuracy and reduce computational load, a three-dimensional region of interest encompassing the neuron of interest was cropped prior to filament tracing. The filament thinnest diameter was set to 0.7 µm, and no maximum filament diameter constraint was applied.

Initial filament seed points and segment assignments were generated automatically and refined through an iterative machine learning-based process involving repeated adjustment of seed placement to improve filament continuity and segmentation accuracy. Disconnected or spurious segments were removed, and missing branches were added manually where necessary to ensure anatomically accurate reconstruction of neuronal processes.

Morphological parameters were extracted directly from Imaris, including Sholl intersections, area under the curve. Sholl analysis was performed using concentric spheres centered on the soma with a fixed radial step size of 10.0 µm, allowing quantification of dendritic complexity as a function of distance from the soma. The area under the Sholl curve (AUC) was calculated by integrating the number of dendritic intersections across radial distance from the soma for each neuron.

Quantitative outputs were automatically exported via ImarisXT and processed in MATLAB for statistical analysis and figure generation. Final datasets were compiled in Microsoft Excel prior to statistical testing.

## Acknowledgments

This work was supported by NINDS award R00NS121399 (QAN), the Esther A. & Joseph Klingenstein Fund (QAN), the Vanderbilt School of Medicine Dean’s Office (LG), and the Cellular, Biochemical, and Molecular Sciences Training Program through the National Institute of General Medical Sciences of the National Institutes of Health T32GM137793 (MD). The authors would like to thank R. Nicoll, D. Debanne, M. Alvarez-Saavedra, D. Bartolome-Martin, K. Zavalin, B. Brown and E. Kavalali for helpful discussions, D. Hartmann for comments on the manuscript, R. Sando lab for assistance with confocal imaging, J. Grewal for assistance with immunostaining and N. Turhan and J. Sandoval for technical assistance. We also thank O. Kovtun and T. Vadakkan from the Vanderbilt Cell Imaging Shared Resource (CISR) Core (supported by NIH grants CA68485, DK20593, DK58404, DK59637, EY08126 and S10OD021630) for their assistance with confocal imaging and Imaris software analysis.

## Data Availability

All data supporting the results presented in the manuscript are included in the figures.

