## Supplementary Figures and Tables for "Distinct electrophysiological signatures reveal neuronal heterogeneity in the mouse fasciola cinereum"

[5] Lead contact

**This file includes:**

Appendix Figures 1 to 6 and Tables 1-2

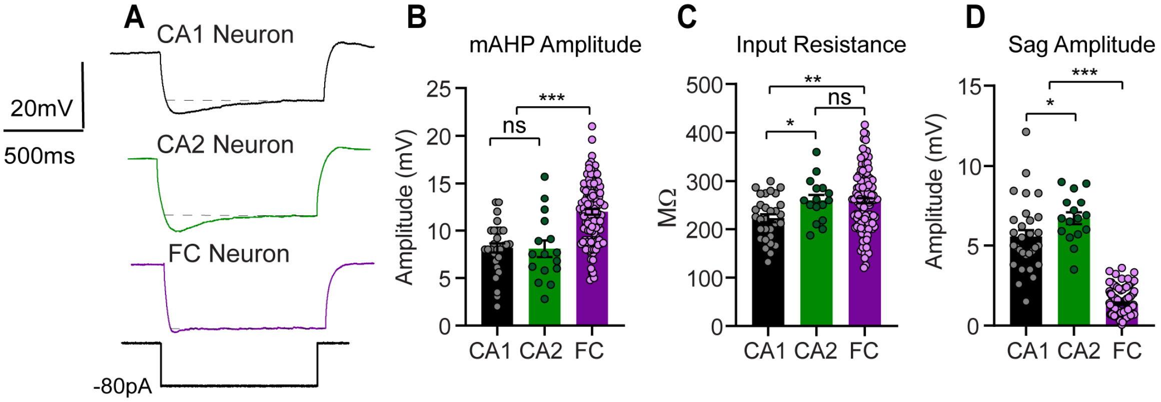


**Figure A1. Intrinsic membrane properties distinguish FC neurons from CA1 and CA2 neurons.**

(A) Representative voltage responses of CA1, CA2, and FC neurons to hyperpolarizing current injection (−80 pA), illustrating differences in subthreshold membrane properties and sag responses. (B) Quantification of medium afterhyperpolarization (mAHP; CA1 PNs= 8.1 ± 0.5 mV; CA2 PNs= 8 ± 0.9 mV; FC= 12 ± 0.3 mV) amplitude showing increased mAHP in FC neurons compared with CA1 and CA2. (C) Input resistance measured across neurons, showing higher input resistance in CA2 and FC neurons compared with CA1 (CA1 PNs= 222 ± 8.2 MΩ; CA2 PNs= 260.6 ± 11.2 MΩ; FC= 259.3 ± 5.8 MΩ). (D) Quantification of hyperpolarization-induced sag amplitude, demonstrating reduced sag in FC neurons relative to CA1 and CA2 (CA1 PNs= 5.5 ± 0.4 mV; CA2 PNs= 6.7 ± 0.4 mV; FC= 1.4 ± 0.1 mV). Data are presented as mean ± SEM; statistical comparisons were performed using one-way ANOVA followed by post hoc multiple comparisons (*$p < 0.05$, **$p < 0.01$, ***$p < 0.001$; ns, not significant). CA1 (n=30 cells/N=9 animals); CA2 (n=15/N 5); FC (n=117/N= 35).

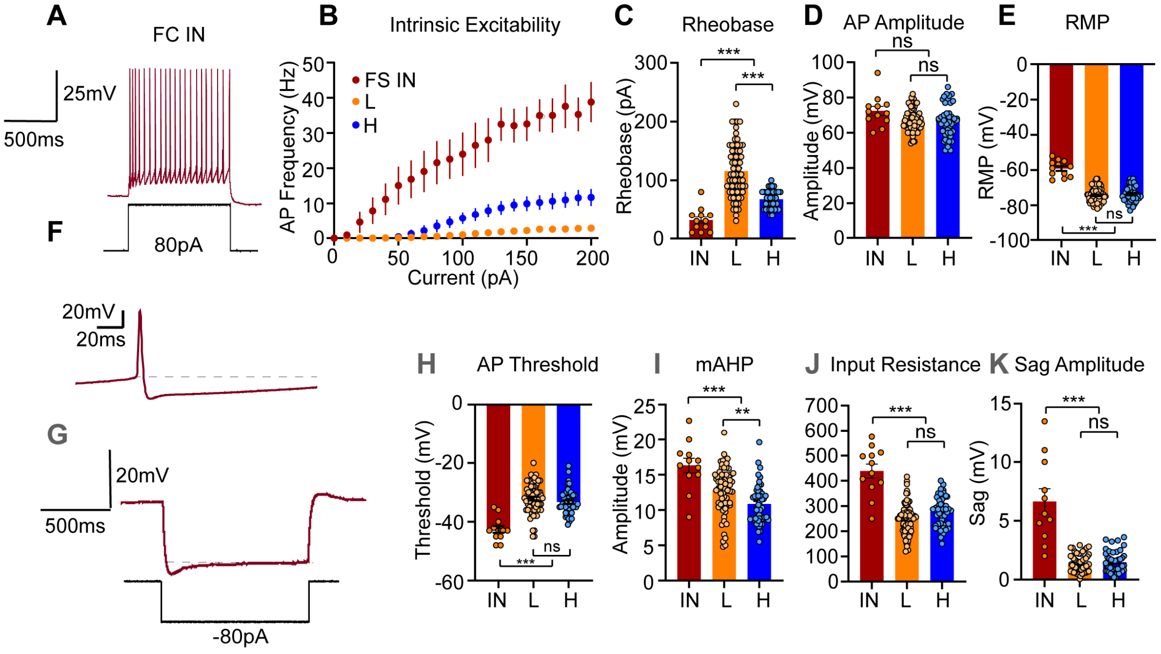


**Figure A2. Electrophysiological comparison of FC principal neurons and interneurons.**

(A) Representative voltage responses of a fast-spiking interneuron (IN) to depolarizing current injection, illustrating high-frequency firing. Scale bars: 25 mV, 500 ms. (B) Input–output relationship showing action potential (AP) firing frequency as a function of injected current for interneurons (IN) and FC principal neurons grouped by excitability (low, L; high, H). Data are presented as mean ± SEM. (C–E) Quantification of intrinsic electrophysiological properties, including rheobase (C; IN= 31.6 ± 5.8 pA; L= 115.7 ± 5.4 pA; H= 67.7 ± 2.4 pA), AP amplitude (D; IN= 72.4 ± 2.5 mV; L= 67.2 ± 0.7 mV; H= 66.4 ± 1.4 mV), and RMP (E; IN= -59 ± 1.3 mV; L= -74.3 ± 0.5 mV; H= -73.7 ± 0.7 mV). (F) Representative single action potential waveform recorded from an interneuron. Scale bars: 20 mV, 20 ms. (G) Representative voltage responses to hyperpolarizing current injection illustrating membrane properties, including sag. Scale bars: 20 mV, 500 ms. (H–K) Quantification of additional intrinsic properties, including AP threshold (H; IN= -42 ± 1.1 mV; L= -32.6 ± 0.3 mV; H= -33 ± 0.6 mV), medium afterhyperpolarization (mAHP; I; IN= 16.3 ± 1 mV; L= 12.84 ± 0.4 mV; H= 10.8 ± 0.4 mV), input resistance (J; IN= 438.4 ± 27.7 MΩ; L= 247.5 ± 7.4 MΩ; H= 277.7 ± 8.7 MΩ), and sag amplitude (K; IN= 6.6 ± 1 mV; L= 1.25 ± 0.1 mV; H= 1.5 ± 0.1 mV). Individual data points represent single cells. Statistical comparisons were performed using one-way ANOVA followed by appropriate post hoc tests (*p < 0.05, **p < 0.01, ***p < 0.001; ns, not significant). IN (n=12 cells/N=10 animals); L (n=71/N=30); H (n=46/ N=20).

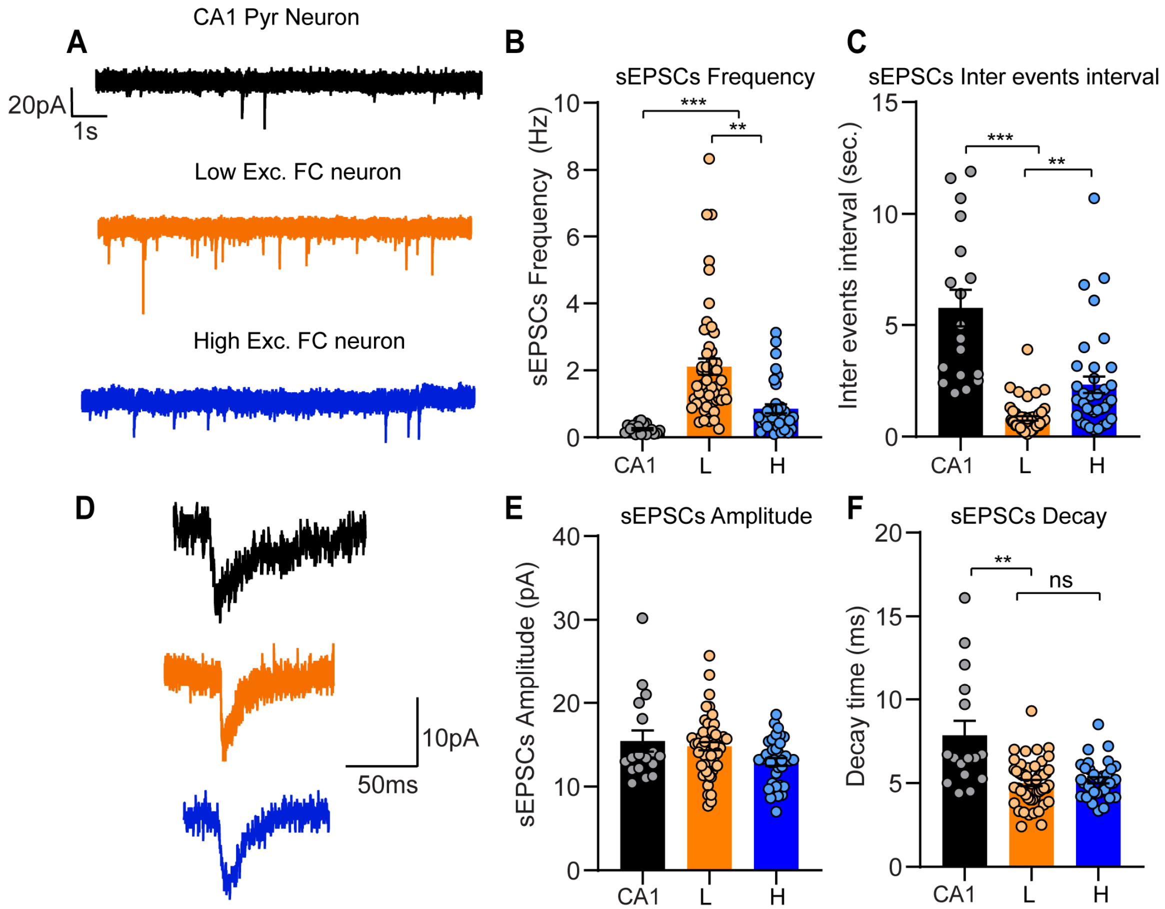


**Figure A3. FC neurons receive enhanced excitatory synaptic input with variability across the population.**

(A) Representative voltage-clamp recordings of spontaneous excitatory postsynaptic currents (sEPSCs) from CA1 neurons and FC neurons spanning the range of intrinsic excitability. (B) Quantification of sEPSC frequency showing increased event frequency in FC neurons compared with CA1. Within the FC population, neurons at the lower end of the excitability range exhibit higher sEPSC frequency than more excitable neurons (CA1 PNs= 0.24 ± 0.03 Hz; L= 2.1 ± 0.2 Hz.; H= 0,85 ± 0.1 Hz). (C) Inter-event interval analysis confirming differences in synaptic event frequency across groups (CA1 PNs= 5.9 ± 0.85 sec.; L= 0.86 ± 0.1 sec.; H= 2.7 ± 0.5 sec.). (D) Representative averaged sEPSC waveforms from CA1 and FC neurons. (E) Quantification of sEPSC amplitude showing no significant differences across groups (CA1 PNs= 15.44 ± 0.1.3 pA; L= 14.82 ± 0.5 pA; H= 12.93 ± 0.5 pA). (F) Quantification of sEPSC decay time indicating a modest difference in synaptic kinetics across conditions (CA1 PNs= 7.8 ± 0.87 ms; L= 5 ± 0.2 ms; H= 5.1 ± 0.3 ms). Data are presented as mean ± SEM; statistical comparisons were performed using one-way ANOVA followed by appropriate post hoc tests (*p < 0.05, **p < 0.01, ***p < 0.001; ns, not significant). CA1 (n=18 cells/N=7 animals); L (n=51/N=25); H (n=35/ N=20).

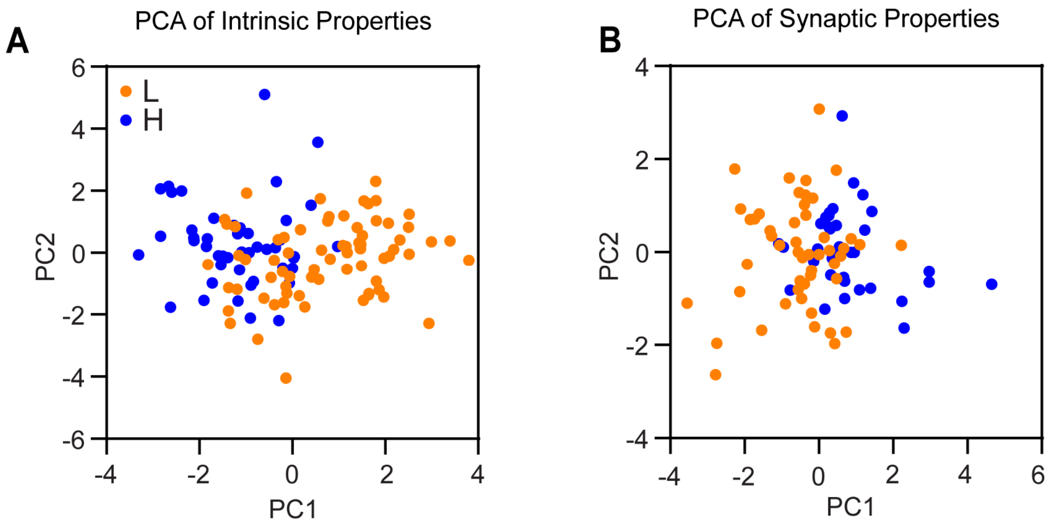


**Figure A4. PCA analysis of intrinsic and synaptic properties.**

(A) Principal component analysis (PCA) incorporating a combined set of intrinsic parameters (Input resistance, rheobase, RMP, sag amplitude, AP amplitude, AP threshold, mAHP, and average I-O spike count). Neurons are color-coded according to relative excitability (low, L; high, H; as denoted in Fig. 2A) to visualize their position along the excitability spectrum. (B) PCA of synaptic properties, including parameters of spontaneous excitatory postsynaptic currents (sEPSCs), showing the distribution of FC neurons based on excitatory synaptic input. Across both intrinsic and synaptic feature spaces, FC neurons exhibit broad and overlapping distributions without clear separable clusters, consistent with a continuum of electrophysiological states. Notably, neurons with lower intrinsic excitability tend to occupy regions associated with higher synaptic input, whereas more excitable neurons are distributed in regions associated with reduced synaptic drive, consistent with an inverse relationship between intrinsic excitability and synaptic input.

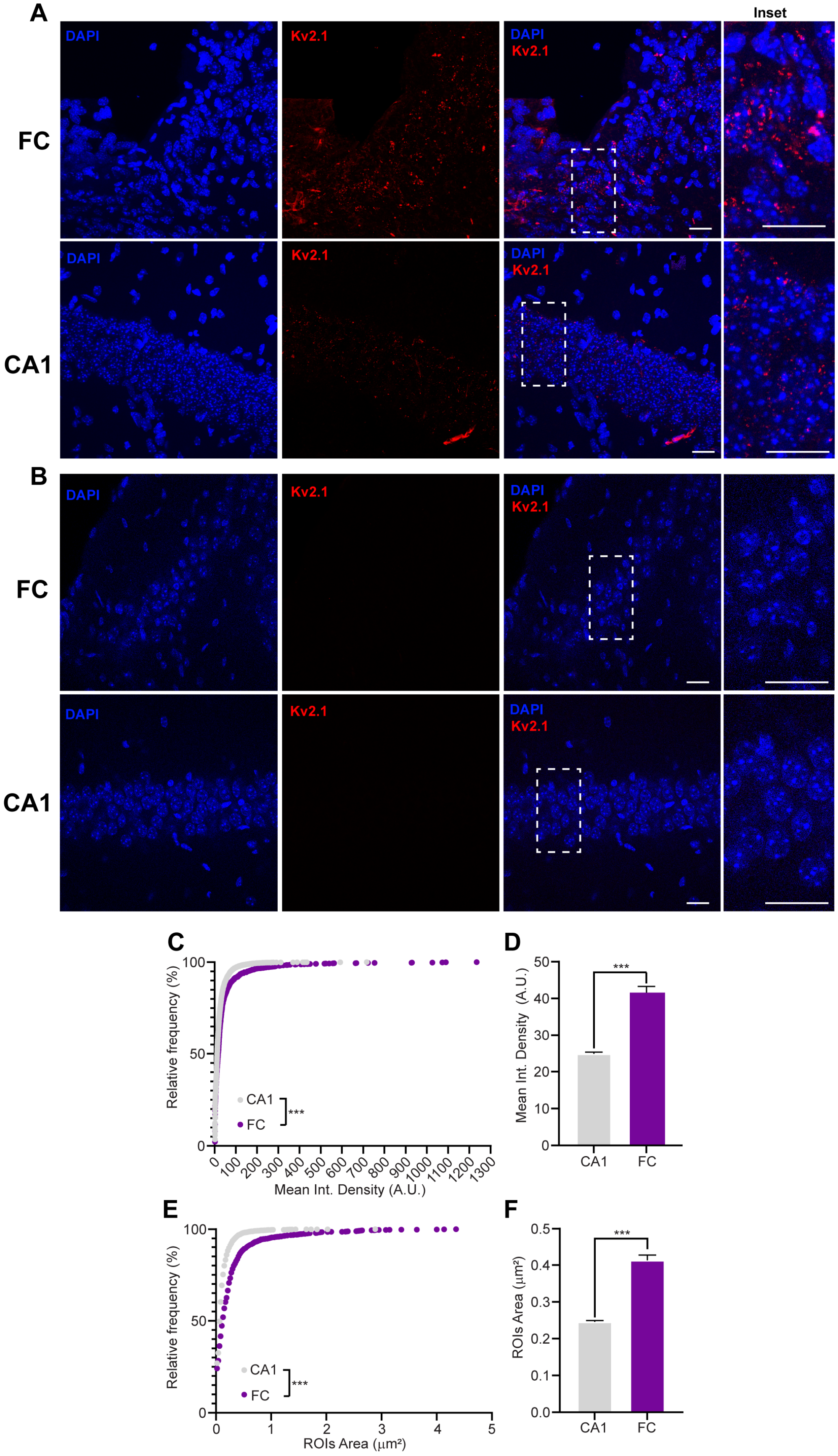
A
Inset
DAPI
Kv2.1
FC
CA1
B
C
Relative frequency (%)
CA1
FC
***
Mean Int. Density (A.U.)
D
Mean Int. Density (A.U.)
E
ROIs Area (µm²)
F
ROIs Area (µm²)

99

**Figure A5. Enriched somatic clustering of Kv2 channels in fasciola cinereum neurons.**

(A) Representative confocal images showing Kv2 immunoreactivity in CA1 and fasciola cinereum (FC) neurons. Kv2 labeling displays a characteristic punctate, somatic clustering pattern, which is more prominent in FC neurons. (B) Representative confocal images showing secondar-only control immunoreactivity in CA1 and FC neurons. (C) Distribution of integrated fluorescence values highlights a right-skewed profile in FC neurons, with a subset of cells exhibiting markedly elevated Kv2 signal. (D) Quantification of integrated Kv2 fluorescence intensity within somatic regions of interest (ROIs) reveals significantly higher signal in FC compared to CA1 neurons (CA1 PN=22 ± 0.4; FC= 40 ± 2.1). (E) Distribution of Kv2-cluster area highlights a markedly elevated area profile in FC compared to CA1. (F) ROI-based analysis of Kv2-positive cluster area shows a significant increase in the total somatic area occupied by Kv2 clusters in FC neurons relative to CA1 (CA1 PN=0.23 ± 0.03; FC= 0.52 ± 0.02). All images were acquired using identical confocal acquisition parameters. Data are presented as mean ± SEM; statistical comparisons were performed using Kolmogorov-Smirnov Normality test (***$p<0.0001$) and Mann-Whitney test (***$p<0.0001$). Scale bars 20μm. Secondary-only control (no primary antibody) shows no detectable signal under identical imaging conditions, confirming the specificity of the immunostaining. CA1 (n=6332 clusters cells/N=3 animals); FC (n=1771/N= 3).

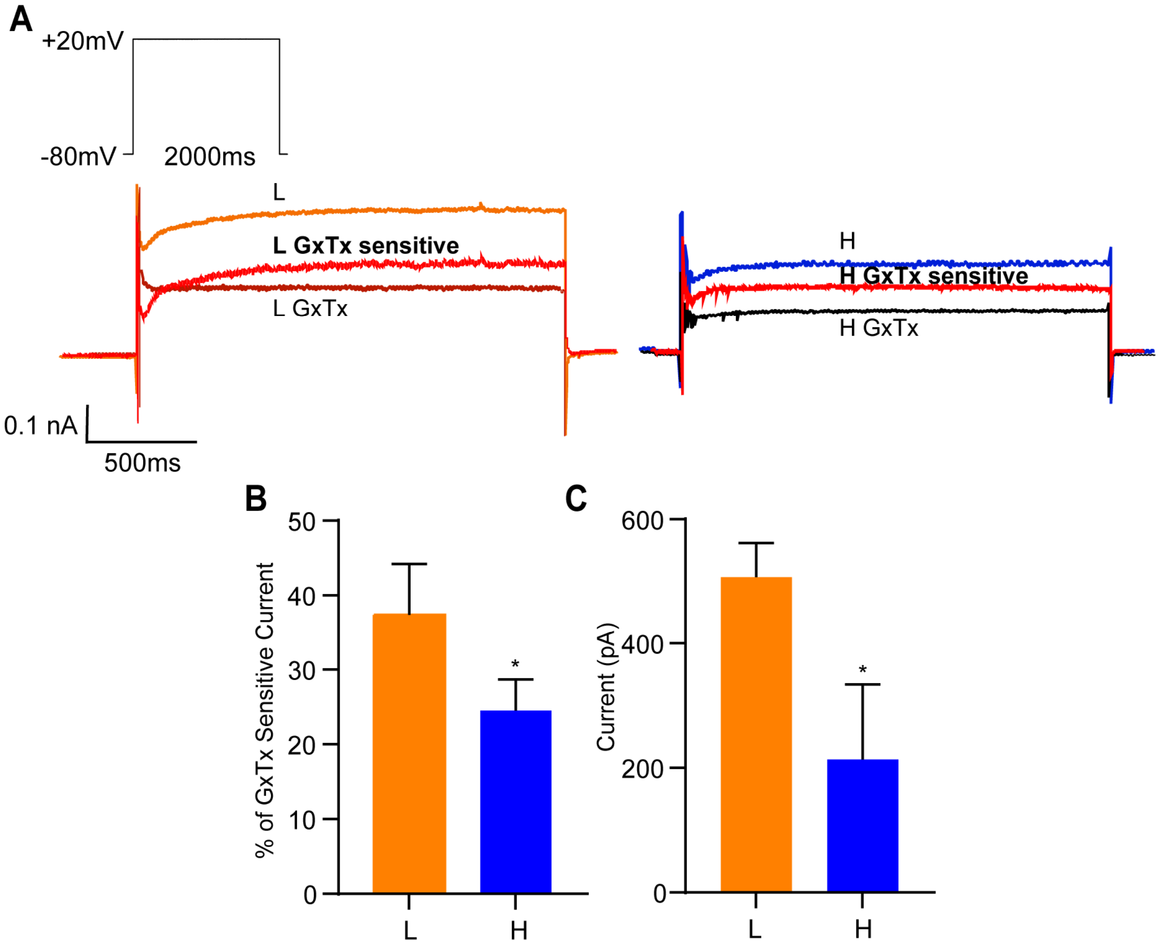


**Figure A6. GxTx-sensitive current varies across FC neurons and is reduced in more excitable cells.**

(A) Representative voltage-clamp recordings from FC neurons illustrating outward potassium currents under control conditions (orange/blue), following application of Guangxitoxin-1E, and the GxTx-sensitive component obtained by point-by-point subtraction (red). Example traces are shown for neurons at the lower (L) and higher (H) ends of the excitability range. (B) Quantification of the GxTx-sensitive current expressed as a percentage of total outward current, showing a reduced contribution in more excitable neurons (L= 37.4 ± 6.8 %; H= 21.7 ± 4.4 %). (C) Absolute amplitude of the GxTx-sensitive current measured at +20 mV, demonstrating a smaller GxTx-sensitive component in more excitable neurons (L= 507 ± 54 pA; H= 209.3 ± 107 pA). Data are presented as mean ± SEM; *$p < 0.05$ by Mann-Whitney test.

**Table A1 (related to figure 3):**

| **Figure 3** | ($I_{Kv2.1}$) -10 mV | 0 mV | 10 mV | 20 mV | 30 mV | 40 mV |
|---|---|---|---|---|---|---|
| FC L (n=32) (pA) | 657.1 ± 36.7 | 990 ± 56 | 1367 ± 78 | 1765 ± 100 | 2177 ± 127 | 2592 ± 153 |
| FC H (n=32) (pA) | 341 ± 41 | 484.3 ± 58 | 653.6 ± 79 | 841.4 ± 103 | 1044 ± 128 | 1244.3 ± 153.8 |

**Table A2 (related to figure 4):**

| **Figure 4** | -10 mV | 0 mV | 10 mV | 20 mV | 30 mV | 40 mV |
|---|---|---|---|---|---|---|
| FC L non-treated (n=9) (pA) | 655.8 ± 106 | 696 ± 141 | 1329 ± 184 | 1756 ± 229 | 2204 ± 281 | 2662 ± 327 |
| FC L GxTx (n=9) (pA) | 506 ± 80 | 722.9 ± 125 | 968 ± 175 | 1223 ± 231 | 1472 ± 275 | 1711 ± 183 |
| $I_{Kv2.1}$ | -10 mV | 0 mV | 10 mV | 20 mV | 30 mV | 40 mV |
| FC H non-treated (n=9) | 400 ± 105 | 538.6 ± 145 | 721.6 ± 188 | 909.6 ± 242 | 1119 ± 294 | 1321 ± 339 |
| FC H GxTx (n=9) (pA) | 297.6 ± 58 | 427.5 ± 88 | 297.6 ± 58 | 685.8 ± 157 | 834 ± 183 | 983 ± 216 |
